# Simulating population pangenomes under coalescent demographic models with MSpangenome

**DOI:** 10.64898/2026.06.29.735168

**Authors:** Lucien Piat, Sukanya Denni, Siegfried Dubois, Benjamin Linard, Ludovic Duvaux

## Abstract

**Motivation:** Pangenome variation graphs (PVGs) are increasingly used to represent genomic diversity, yet there is currently no general framework for generating population pangenomes directly from explicit evolutionary histories. Existing simulators typically focus on individual classes of variation and do not integrate these variations within a genealogy-aware framework driven by explicit demographic histories. As a result, evaluating pangenome methods in realistic population-genetic settings remains challenging, and benchmark datasets with known evolutionary ground truth are scarce.

**Results:** We present `MSpangenome`, a genealogy-aware framework that bridges coalescent population genetic simulations and pangenome graph analyses. The pipeline combines ancestry simulation with `msprime` and a *de novo* graph construction algorithm to generate PVGs directly from simulated genealogies. By explicitly modeling recombination, demographic history and incomplete lineage sorting, `MSpangenome` produces structurally complex pangenomes in which nested and overlapping structural variants emerge naturally from the underlying genealogies, while their evolutionary history and graph topology remain known by construction. This provides a general framework for generating realistic population pangenomes and establishing ground-truth datasets for methodological evaluation. We demonstrate its utility by generating population-scale pangenomes and using them as controlled references to benchmark the widely used graph construction tools, `PGGB` and `Minigraph-Cactus`. Our analyses reveal contrasting performance regimes across levels of sequence diversity, sample sizes and classes of structural variation, highlighting the value of simulation-based benchmarking for identifying reconstruction errors that are hard to detect using empirical datasets alone.

**Availability and implementation:** `MSpangenome` is implemented in Python, fully containerized, freely available at https://forge.inrae.fr/pangepop/MSpangepop and mirrored at https://github.com/inrae/MSpangepop.

## Introduction

Historically, genetic diversity has been described primarily using single-nucleotide polymorphisms (SNPs) and microsatellites. However, genomic variation also includes structural variants (SV), generally defined as mutations larger than 50 bp. These variants can alter chromosome structure on medium to large scales and can contribute significantly to the genetic diversity of populations and may have major phenotypic effects (Catlin et al., 2025; Oliver et al., 2013). Despite their importance, SVs have long been overlooked in genomic analyses and remain poorly modeled in evolutionary frameworks. On the one hand, genetic diversity simulators grounded in rigorous theory do not explicitly incorporate SVs (Baumdicker et al., 2021; Haller and Messer, 2017). On the other hand, simulators specifically designed to generate SVs like VISOR or VACsim (Bolognini et al., 2020; Ding et al., 2026) rely on simplified evolutionary assumptions, including basic demographic models (e.g. star-like phylogenies) and/or constrained mutation schemes (e.g. absence of nested or overlapping SVs, where mutations can occur within the sequence introduced by previous variants). Together, these limitations reveal a critical gap in our ability to simulate SVs under realistic evolutionary models, thereby limiting benchmarks and downstream biological inference.

Recently, pangenomics has emerged as a powerful conceptual framework to better comprehend genome structural complexity and the contribution of structural variants (SVs) to phenotypic variation and genome evolution (Bocs et al., 2025). Initially developed for prokaryotes, it has now been extended to the complexity of eukaryote genomes, where it can describe the full set of genetic sequences within a population or species by integrating both SNPs and SVs. Since the early development of pangenomics, several formalisms, including de Bruijn graphs (Leggett et al., 2013), have been proposed to represent pangenomes, but they remain limited in their ability to capture large-scale rearrangements and repetitive regions (Andreace et al., 2023), especially in structurally complex and repeat-rich genomic regions. In this context, pangenome variation graphs (PVGs) provide an alternative representation that trades compression for improved handling of SVs and easier biological interpretation (Fig. 1). Briefly, paths through the graph describe individual haplotypes, separating shared regions from haplotype-specific variants and thereby facilitating SV detection (Groza et al., 2024). Due to the increasing availability of high-quality whole genome assemblies, tools such as `PGGB` (Garrison et al., 2024) and `MinigraphCactus` (Hickey et al., 2023) allow the construction of PVGs. However, methodological evaluation and experience with these recent approaches remain limited, and robust frameworks for graph quality benchmarking are still scarce (see Kopalli et al., 2025). Moreover, the lack of simulators capable of generating realistic synthetic pangenomes with known PVG or alignment ground truth prevents objective evaluation. Together, these limitations constitute a major methodological bottleneck, further compounded by reproducibility concerns.

**Fig. 1.**
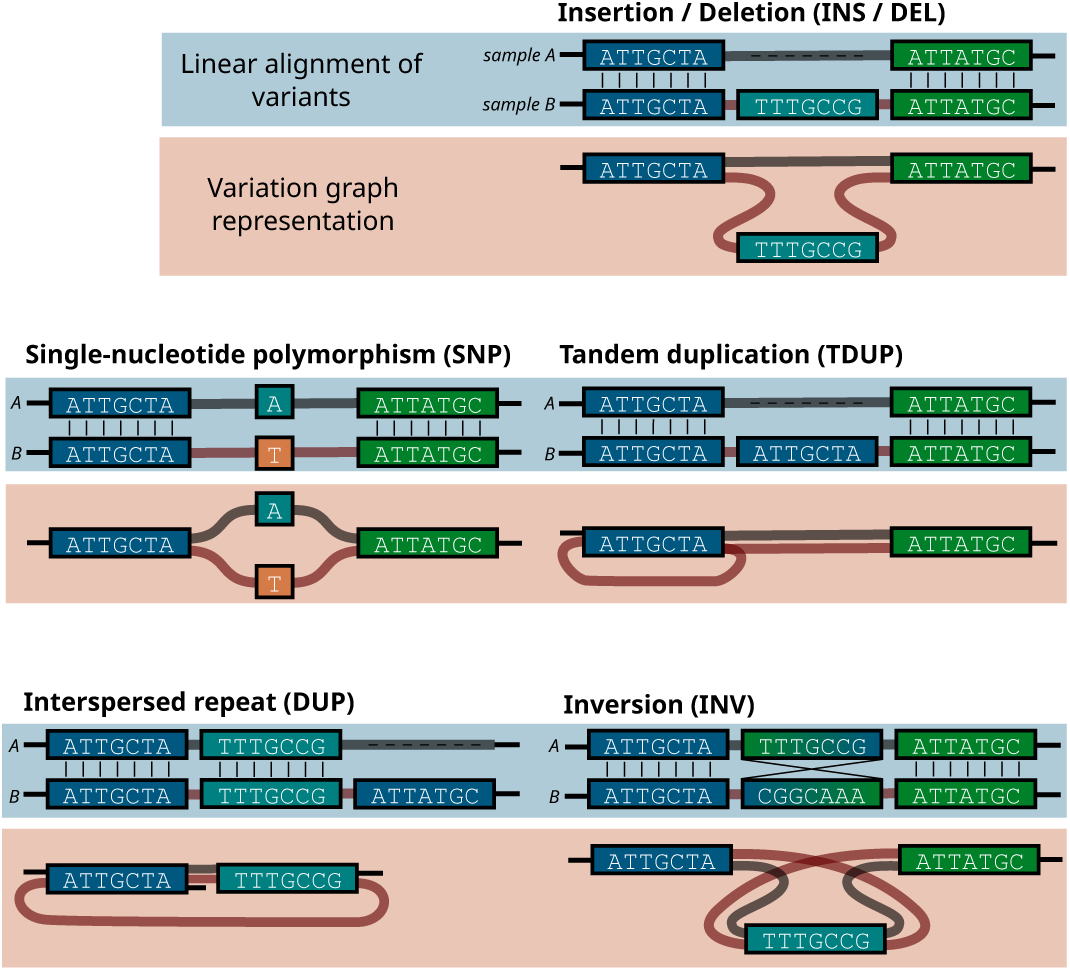
Genomic structural variations represented in linear and graph formats. Two haplotypes, sample *A* and *B*, are shown as either a conventional linear alignment (upper panels) or a variation graph (lower red panels). Colored blocks represent homologous genomic segments. In the graph representation, each haplotype corresponds to a distinct path through the graph. Structural variants are represented as alternative paths, while repeated traversals of graph nodes generate tandem and interspersed duplications. Edges connecting identical node extremities (5, *−* 5, or 3, *−* 3,) indicate a strand reversal and therefore traversal of the corresponding sequence in reverse-complement orientation. The inversion example illustrates such a reversal.

To address this critical gap, we developed `MSpangenome`, a simulator that generates biologically realistic synthetic pangenomes with fully controlled structure by integrating coalescent-based simulations of demo-genetic history with a novel algorithm for PVG generation. This tool provides a controlled reference framework for the rigorous benchmarking of pangenomic methods, enabling the reproducible generation of standardized evaluation datasets and laying the foundation for robust and precise validation of structural variation at scale. We describe the pipeline in detail, illustrate its application with a toy bacterial genome, and demonstrate its utility through a comparative evaluation of existing graph construction approaches.

## Results

### Overview

Population pangenome simulation requires addressing several interrelated challenges: realistic demographic modeling, recombination-driven genealogical heterogeneity, the generation of complex structural variants and the direct production of standards-compliant graph-based genome representations (i.e. GFA). `MSpangenome` addresses these requirements through a five-step workflow comprising ancestry simulation, genealogy traversal, variant generation, graph construction and graph compaction (Fig. 2). The workflow is implemented in Python using a modular object-oriented architecture, orchestrated by Snakemake (Mölder et al., 2021), and fully containerized to ensure reproducibility across computing environments.

**Fig. 2.**
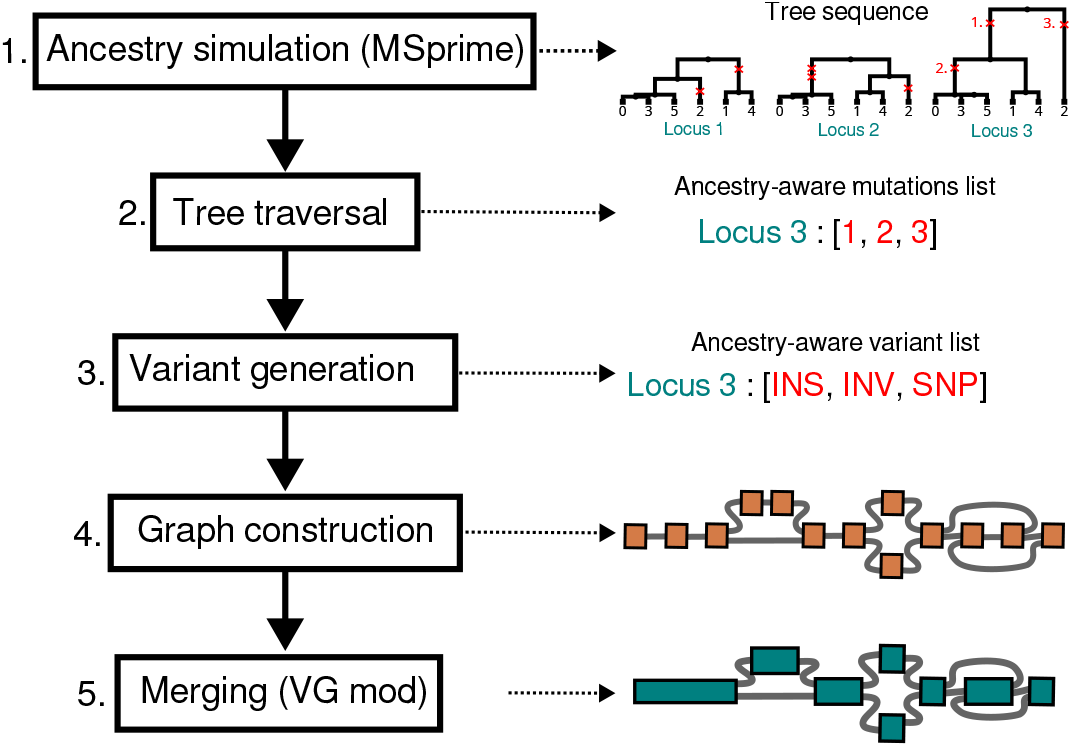
`MSpangenome` workflow overview. The pipeline comprises five sequential stages. **(1) Ancestry simulation:** `msprime` generates a tree-sequence containing multiple local genealogies arising from recombination; mutations (red marks) are placed along branches under a Poisson mutation process with rate *µ*. **(2) Tree traversal:** Preorder traversal of each local genealogy extracts mutations in chronological order (oldest first), yielding a mutation list record per locus. **(3) Variant generation:** Each mutation is assigned a variant type (SNP, DEL, INS, INV, DUP) drawn from user-specified distributions. **(4) Graph construction:** Variants are sequentially applied to build an expanded pangenome graph where each node represents a single nucleotide. **(5) Node compaction:** Linear single nucleotide node chains are concatenated into single nodes, reducing graph size while preserving topology.

### Coalescent-based ancestry simulation

To generate biologically realistic genealogies, `MSpangenome` leverages `msprime` (Baumdicker et al., 2021), a coalescent simulator that efficiently models recombination using a treesequence, a data structure that compactly stores the succession of local genealogies along the genome (Tsambos et al., 2023). Recombination naturally partitions the genome into loci, each characterized by its own local genealogy. Neighboring loci retain correlated genealogies, giving rise to linkage disequilibrium, a fundamental property of real genomes that cannot be captured by single-tree models. At the same time, differences in the shape and age of local genealogies also result in varying densities of mutations across loci.

Users specify two categories of parameters: demographic specifications (e.g. population sizes *N*_*e*_, split times, migration rates) and genomic parameters (e.g. mutation rate *µ*, recombination rate *r*, genome length *L*). These parameters can be combined into composite quantities such as the population-scale mutation rate *θ* = 4*N*_*e*_*µ*, which captures the expected level of neutral polymorphism, and the population-scale recombination rate *ρ* = 4*N*_*e*_*rL*, which determines the number of independent genealogies along a chromosome. Complex population structures can be modeled using JSON-formatted demographic scenarios, either defined using the `demes` standard (Gower et al., 2022) or imported from the `stdpopsim` catalog (Adrion et al., 2020; Lauterbur et al., 2023); (see sup Tab. 4 and sup Tab. 5 for the full list of tunable parameters). To maintain computational tractability, whole-genome simulations are parallelized at the chromosome level, naturally reflecting the independent assortment of chromosomes during meiosis.

### Complex variant generation

Once genealogies are simulated, `msprime` places mutation events along the branches of each local tree. We designed an algorithm that uses msprime’s tree collections to extract these mutations via a preorder traversal of each genealogy, yielding a chronologically ordered list where older mutations precede more recent ones, which is critical for the correct handling of nested variants.

Each mutation is then converted into a genetic variant by assigning it a type (insertion, deletion, inversion, tandem duplication, or SNP), a position, and a size—drawn from userdefined or default distributions. The SV categories implemented in `MSpangenome` were inspired by those supported in VI-SOR (Bolognini et al., 2020), while extending their use to mutations evolving along explicit local genealogies. A central challenge in handling these variants is maintaining coordinate consistency across the mutation list. When a variant of size *k* is introduced at position *p*, the coordinate system must be adjusted for all subsequent mutations: deletions shrink the available interval, removing potential mutation targets, while insertions expand it, creating new substrate for future variants. This dynamic propagation mechanism naturally produces nested variants—for instance, a SNP occurring within a previously inserted sequence—while maintaining topological consistency throughout the graph. The resulting coordinate updates, however, translate into a significant computational cost. Because each structural modification must be propagated to all subsequent mutations in the lineage, processing a lineage containing *m*_*l*_ mutations has a worstcase time complexity of 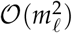 per locus. At the genome scale, worst-case time complexity is therefore bounded by *O* (*m*^2^), where *m* is the total number of mutations. In practice, parallelization across loci substantially mitigates this cost.

### Pangenome graph construction

The enriched mutation list, now containing both genealogical context and variant specifications, serves as a set of instructions for the graph construction algorithm. The implementation follows an object-oriented design built around five core classes: `Node` (representing a single nucleotide), `Edge` (a bidirectional connection between node extremities), `Path` (an ordered edge list representing a haplotype), `Graph` (subgraph for one locus), and `GraphEnsemble` (chromosome-level collection of subgraphs).

We adopt an expanded representation (Garrison et al., 2024; Guarracino et al., 2022) in which each node contains exactly one base. This fine-grained representation simplifies variant application logic at the cost of increased node count. Crucially, the graph is bidirected: edges connect the 5, or 3, extremities of nodes rather than the nodes themselves, thereby encoding sequence orientation, analogous to the formalism used for compacted de Bruijn graphs (Chikhi et al., 2016). This representation naturally accommodates inversions, which traverse nodes in reverse orientation, and tandem duplications, which revisit existing nodes, without requiring artificial node duplication (see Fig. 1).

Graph construction proceeds in two phases (Fig. 3). During initialization, the ancestral sequence (i.e. any sequence of interest chosen by the user) is converted into a linear chain of nodes connected by forward-oriented edges—a structure we term *string of pearls*. At this stage, all haplotypes share a single identical path through the graph (Fig. 3B). Variants are then applied sequentially, following the chronological order established during tree traversal. For each variant, the `Graph` class identifies the affected genomic interval, translates positions into node identifiers, and delegates the topological modification to the appropriate `Path` objects. Insertions create new nodes and redirect affected paths through them (Fig. 3C); SNPs create alternative nodes at specific positions (Fig. 3D); inversions alter the connectivity, or graph incidence, of edges to nodes extremities without adding nodes, creating characteristic loop structures (Fig. 3E). Because mutation positions are expressed relative to the state of each path at the time they occur, variants arising within previously inserted sequences are automatically nested. A fallback mechanism ensures robustness: if a variant would compromise graph connectivity, the operation is rolled back and logged for inspection. Per-subgraph complexity is *O ·* (*n* (*L*_locus_ + *V*_locus_)), where *n* is the number of individuals, *L*_locus_ the locus length, and *V*_locus_ the variant count.

**Fig. 3.**
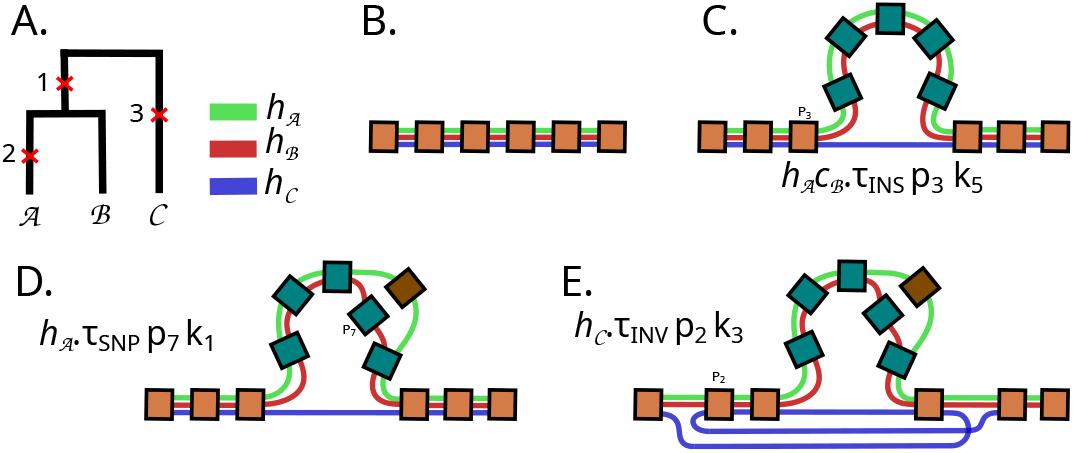
Incremental graph construction through sequential variant application. In this example, we describe the procession of a single locus for three haplotypes (*h*_*A*_, *h*_*B*_, *h*_*C*_), associated to green, red and blue colours respectively. **(A)** Preorder tree traversal dictates the processing order of generated mutations 1 to 3, that occured at different branch position in the tree. Each mutation is encoded as an atomic graph edit operation *h*.*τ*_type_ *p*_*i*_*k*_*j*_, where *h* are the haplotypes involved, *τ*_type_ the variant type, *p*_*i*_ the relative position, and *k*_*j*_ the size. **(B)** Initial state: all haplotypes share a single ancestral path (string of pearls). **(C)** An ancestral insertion (mutation 1) of size *k*_5_ at position *p*_3_ creates new nodes (turquoise) and modifies paths *h*_*A*_ and *h*_*B*_. **(D)** A point mutation (mutation 2) affects only lineage *A*, creating a SNP relative to lineage *B*. Position *p*_7_ is relative to the previously modified path, producing a nested variant. **(E)** An inversion (mutation 3) affects lineage *C*. No new nodes are created; only edge orientations change, forming a topological loop (blue path) through existing nodes.

### Output generation

To limit memory footprint, subgraphs are initially stored in temporary files rather than kept in memory. Chromosome-level assembly then proceeds by sequentially concatenating adjacent locus graphs, connecting the terminal node of each subgraph to the first node of the next. This modular strategy requires that locus boundaries remain mutation-free to guarantee proper connectivity.

The expanded graph is serialized to GFA format using parallelized I/O operations. Node definitions (S-lines) and path/edge reconstruction (P- and L-lines) are handled by separate processes with buffered writes, and final node identifiers are assigned only at export to reduce memory consumption during processing. A companion FASTA file containing the simulated haplotype sequences is generated simultaneously.

Because the expanded graph representation produces one node per nucleotide, the raw GFA files are large. Post-processing compression—termed *unchopping*—merges linear chains of nodes into single nodes (Guarracino et al., 2022) (Fig. 2.5). We perform this step using `vg mod -unchop` (Garrison et al., 2018), which substantially reduces both file size and node count while fully preserving graph topology. Given the sheer volume of single-nucleotide nodes that must be traversed and merged, this indispensable reduction step constitutes the most time-consuming phase of the workflow.

**Fig. 4.**
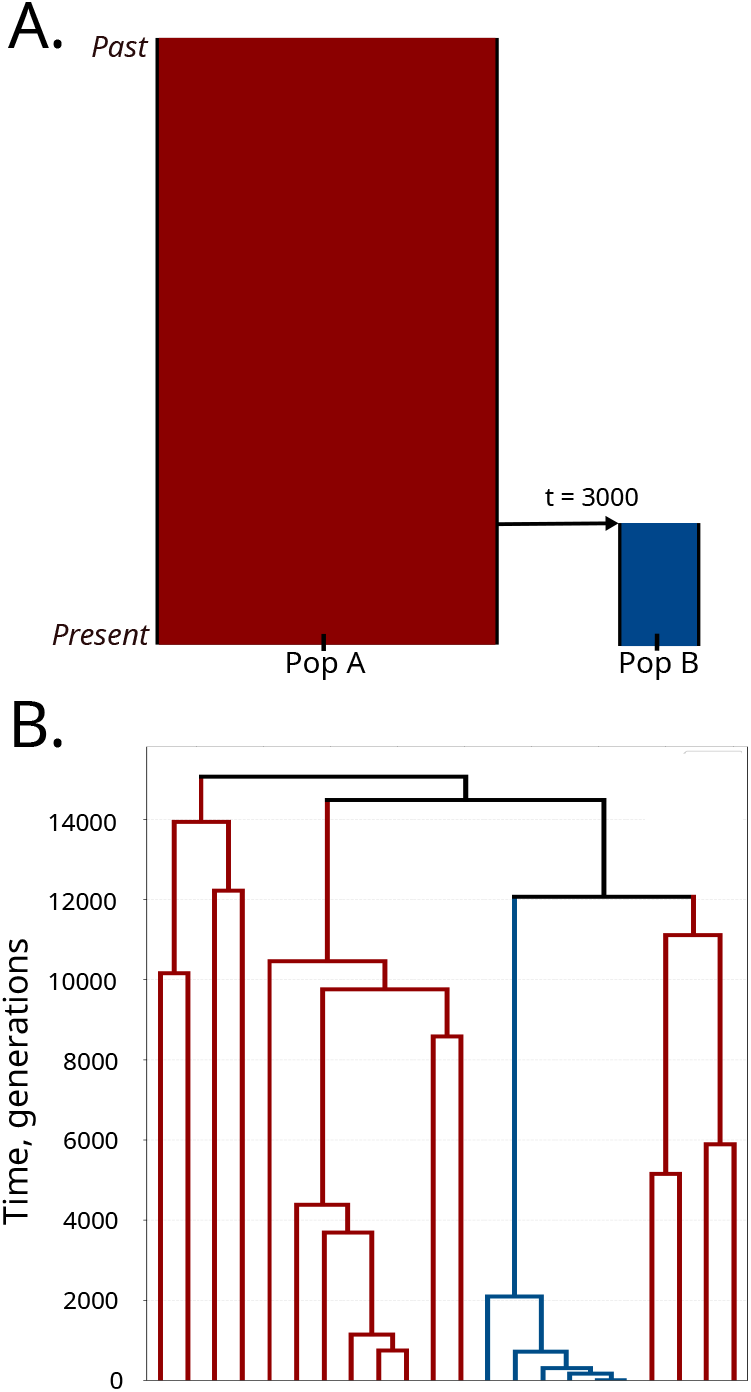
Demographic model and average genealogy of the simulated *S. aureus* pangenome. **(A)** Demographic scenario: Population 2 (blue) diverges from Population 1 (red) at *T* = 3,000 generations. **(B)** STEAC consensus tree computed across all 240 loci.

**Fig. 5.**
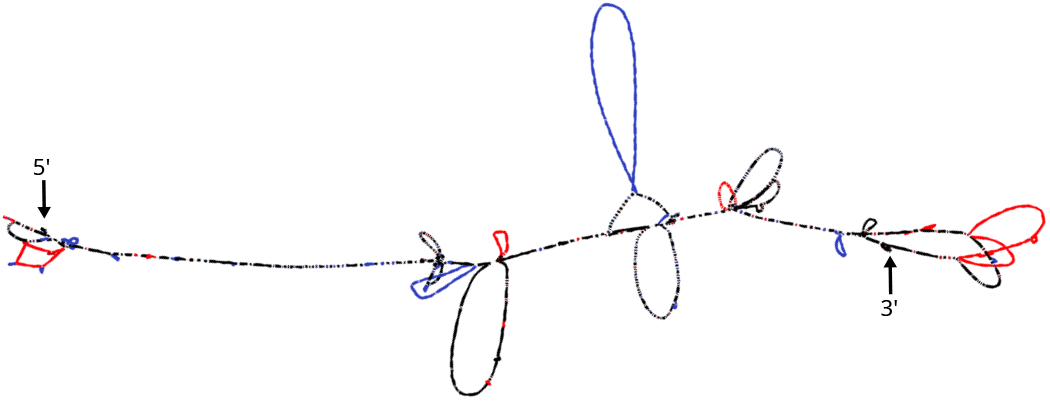
Bandage visualization of the simulated pangenome graph. Nodes colored by population specificity: black (core genome shared by both populations), red (Population 1-specific), blue (Population 2-specific).

### Other functionalities

Beyond graph generation, `MSpangenome` provides built-in analysis tools to facilitate interpretation of simulation results. Visualizing tree-sequences containing hundreds of local genealogies is impractical, so we implemented a module based on STEAC (Species Tree Estimation using Average Coalescence) (Liu et al., 2009). This approach computes pairwise average coalescence times across all loci, producing a genetic distance matrix from which a consensus tree summarizing overall evolutionary history can be reconstructed.

`MSpangenome` also generates comprehensive log files recording every introduced mutation along with its type, time of occurrence, locus assignment, and affected lineages. By avoiding formats tied to a single linear reference (e.g. VCF or BED), this registry serves as ground truth for benchmarking the sensitivity and precision of variant detection tools operating on pangenome graphs. These supplementary functionalities are described in more detail in the documentation wiki of the `MSpangenome` repository.

### Proof of concept: the *Staphylococcus aureus* pangenome

To illustrate the capabilities of `MSpangenome` in a biologically meaningful setting, we simulated a pangenome based on the *Staphylococcus aureus* genome (~3 Mb) (Gillaspy et al., 2014). Although bacterial recombination patterns differ substantially from those of eukaryotes, its relatively compact genome provides a computationally efficient test case that can be rapidly reproduced, while retaining sufficient complexity to exercise all components of the pipeline.

The demographic scenario models a simple population divergence (Fig. 4A), representative of a bacterial subculture event in which a subset of individuals founds a new population. Population 2 derives from Population 1 at time *T* = 3,000 generations ago, with a ten-fold reduction in effective size (*N*_2_ = 500 versus *N*_1_ = 5,000). This asymmetry is expected to produce distinct coalescent patterns between populations. The complete set of simulation parameters is provided in Table 1.

**Table 1.**
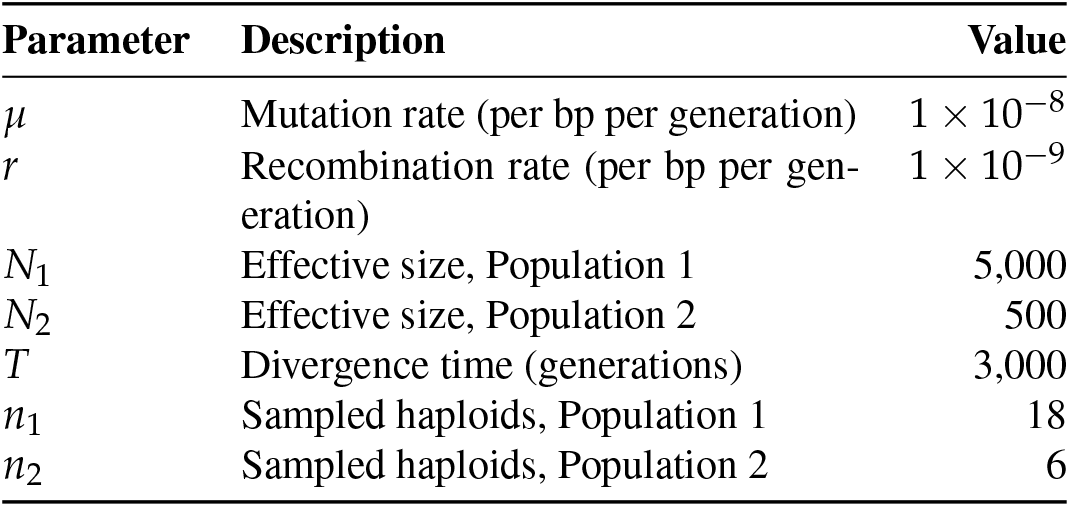
Simulation parameters for the *S. aureus* pangenome.

| Parameter | Description | Value |
| --- | --- | --- |
| $\mu$ | Mutation rate (per bp per generation) | $1 \times 10^{-8}$ |
| $r$ | Recombination rate (per bp per generation) | $1 \times 10^{-9}$ |
| $N_1$ | Effective size, Population 1 | 5,000 |
| $N_2$ | Effective size, Population 2 | 500 |
| $T$ | Divergence time (generations) | 3,000 |
| $n_1$ | Sampled haploids, Population 1 | 18 |
| $n_2$ | Sampled haploids, Population 2 | 6 |

The simulation produced a pangenome partitioned into 240 local genealogies (loci) averaging 12.5 kb in length, carrying a total of 2,603 mutations, corresponding to an average of 11 mutations per locus.

To summarize this complex genealogical structure, we applied the STEAC method (Fig. 4B). As expected from coalescent theory, Population 2 lineages (blue) coalesce substantially faster than those of Population 1 (red), reflecting the accelerated genetic drift imposed by its smaller effective size. Meanwhile, Population 1 appears paraphyletic in the consensus tree—a signature of incomplete lineage sorting that arises when ancestral polymorphism has not yet been lost through genetic drift.

Because the simulated pangenome remains computationally tractable, we visualized the complete graph using Bandage (Wick et al., 2015) (Fig. 5). The resulting topology is broadly consistent with empirical bacterial pangenome graphs (Gautreau et al., 2020), comprising a shared core backbone (black nodes) corresponding to the persistent genome, interspersed with population-specific branching regions and variation bubbles (red and blue nodes) that represent the shell and cloud accessory genomes. Notably, the graph contains nested variants and locally dense substructures, demonstrating `MSpangenome`’s capacity to generate complex structural variation patterns comparable to those observed in empirical pangenomes.

### Comparative analysis of pangenome graph construction tools

Two tools currently dominate eukaryotic PVG construction: `PGGB` (Garrison et al., 2024) and `Minigraph-Cactus` (MC) (Hickey et al., 2023). These tools employ fundamentally different algorithmic strategies. `PGGB` adopts a referencefree approach: it performs all-versus-all alignment using `wfmash` (Guarracino et al., 2024), from which `seqwish` (Garrison and Guarracino, 2022) constructs directly an initial variation graph encompassing all haplotypes; a second smoothing step with `smoothxg` (Garrison et al., 2024) simultaneously simplifies topology and introduces base-level variations through partial order alignment (POA). In contrast, MC employs a reference-guided strategy, combining `Minigraph` to build a structural variant skeleton through iterative alignment starting from a selected reference, and then performs fine-grained sequence alignment and integrates smaller variants. These contrasting approaches can produce markedly different graph topologies from the same input data, yet their relative performance has remained difficult to assess due to the lack of datasets with known ground-truth structure.

Although both tools are widely adopted, their ongoing active development can inherently affect result stability and topological quality. To systematically evaluate their reconstruction fidelity under varying conditions, we leveraged the fully known evolutionary histories generated by `MSpangenome` as ground-truth references. This strategy provides a controlled benchmarking framework that concurrently demonstrates the simulator’s practical utility for pangenomic tool assessment.

#### Experimental design

Our benchmarking analysis followed a three-stage experimental design (Fig. 6):

**Fig. 6.**
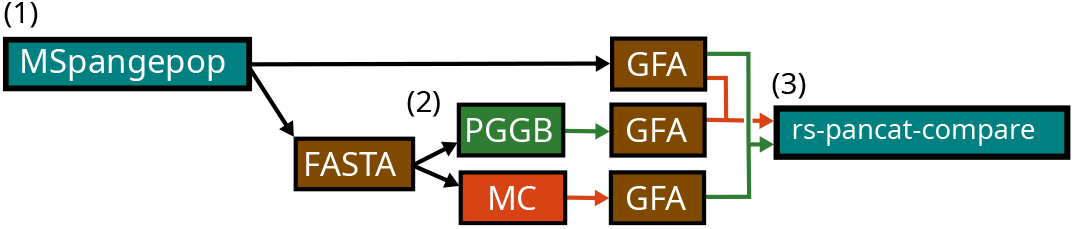
Experimental design for benchmarking pangenome graph construction tools. The workflow comprises three stages: (1) simulation of reference graphs with `MSpangenome` generating 510 datasets (17 conditions *×* 30 replicates); (2) reconstruction from 510 simulated FASTA files sequences using `PGGB` and MC, and (3) quality assessment via Path Edit Distance (PED). Box colors designate benchmarking tools (teal), `PGGB` (green), MC (orange), and intermediate files (brown).

1. **Simulation:** We generated 510 pangenome graphs with `MSpangenome` v0.1.3 based on the *S. aureus* genome, comprising 17 experimental conditions with 30 simulation replicates each (Table 2). All simulations assumed a single constant-size panmictic population (*N*_*e*_ = 5000) and a fixed recombination rate of *r* = 8 *×* 10^*−*9^ per base per generation. Experimental conditions differed with respect to three factors (each varied separately unless otherwise specified): genetic diversity (*θ*), the number of sampled haplotypes (*n*), and the relative proportion of variant types (SNP, DEL, INS, INV, DUP). Variant frequencies followed either a balanced distribution, *D*_0_ = (30%, 20%, 20%, 20%, 10%), or an enriched distribution, *D*_*τ*_, in which one structural variant class (DEL, INS, INV, or DUP) was increased to 55% while all remaining variant classes were set to 11%.
2. **Reconstruction:** From the simulated FASTA sequences, we reconstructed pangenome graphs using both `PGGB` and MC under default parameters. Because MC requires a reference, the first simulated haplotype of each dataset was arbitrarily chosen as the reference, and the full, unfiltered graph was retained to preserve the complete topology. In total, 1,020 graphs were reconstructed.
3. **Comparison:** We quantified reconstruction quality using Path Edit Distance (PED)—the minimum number of elementary operations (node/edge insertions, deletions, or substitutions) required to transform the paths of one graph into another—computed between each simulated reference and its corresponding reconstructions using `rspancat-compare` (Dubois et al., 2025).

**Table 2.** Benchmark conditions. Varying parameters are indicated in bold.

| Condition | $\theta$ | $n$ | SV distribution |
| --- | --- | --- | --- |
| Base level | $10^{-4}$ | 6 | $D_0 = (30\%, 20\%, 20\%, 20\%, 10\%)$ |
| Diversity (Fig. 7) | $\{0, 2 \times 10^{-5}, 2 \times 10^{-4}, 2 \times 10^{-3}, 10^{-2}\}$ | 6 | $D_0$ |
| Variant type (Fig. 8) | $10^{-4}$ | 6 | $\{\mathbf{D_{DEL}}, \mathbf{D_{INS}}, \mathbf{D_{INV}}, \mathbf{D_{DUP}}\}$ |
| Diversity × size (Fig. 9) | $\{10^{-4}, 10^{-3}\}$ | $\{2, 6, 12, 24\}$ | $D_0$ |

This analysis required approximately 4,200 CPU-hours on a high-performance computing cluster, corresponding to the CO_2_ equivalent of a 70 km journey by combustion-engine vehicle (evaluation provided by the Génotoul HPCC (Berthoud et al., 2020)).

#### Effect of genetic diversity

We first evaluated the impact of population-level genetic diversity, parameterized by *θ* = 4*N*_*e*_*µ*, across five levels: *θ ∈ {*0, 2 *×* 10^*−*5^, 2 *×* 10^*−*4^, 2 *×* 10^*−*3^, 10^*−*2^ *}*. As anticipated from alignment theory, PED exhibits a positive correlation with *θ* (Fig. 7A), reflecting the increasing difficulty of establishing sequence homology under high divergence.

**Fig. 7.**
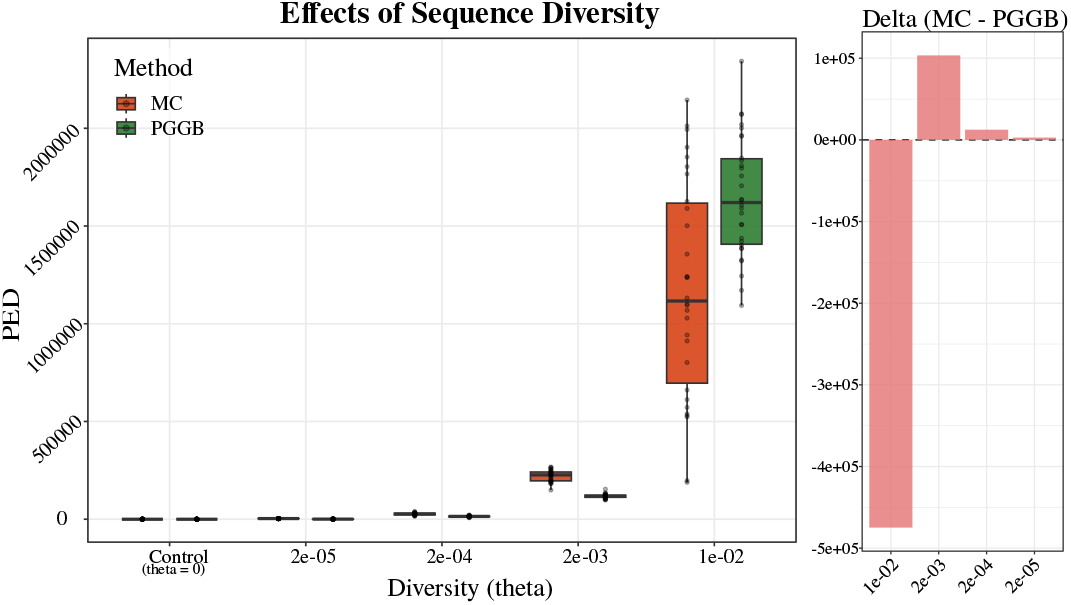
Effect of genetic diversity (*θ*) on graph reconstruction quality. **(A)** PED distribution for MC (orange) and `PGGB` (green) across diversity levels. Significance assessed via paired Wilcoxon test (two-tailed). **(B)** Mean PED difference between methods (Δ = PED_MC_ *−* PED_PGGB_). Positive values indicate superior `PGGB` performance; negative values indicate superior MC performance.

Differences between tools are statistically significant across all diversity levels (paired Wilcoxon test, *p <* 0.05), but the analysis reveals two distinct regimes. For low-to-moderate diversity (*θ ≤* 2*×* 10^*−*3^), PED increases progressively with marginally better performance for `PGGB` (Fig. 7B). A sharp transition occurs at *θ* = 10^*−*2^, a quite high population mutation rate, where PED increases by approximately two orders of magnitude for both methods. This threshold likely marks the saturation limit of alignment heuristics, beyond which mutation density overwhelms the signal used to establish homology.

Interestingly, MC outperforms `PGGB` at this extreme diversity level. However, this advantage is accompanied by a substantially higher variance, suggesting that while MC’s reference-guided anchoring provides better support than `PGGB`’s all-versus-all approach under extreme divergence, reconstruction stability remains precarious in these conditions.

#### Sensitivity to variant types

We next examined whether specific variant classes pose particular challenges for graph reconstruction. We generated graphs in which each structural variant class was alternately enriched to 55% of total variants. Across all conditions, `PGGB` consistently outperforms MC with lower inter-condition variability (Fig. 8).

**Fig. 8.**
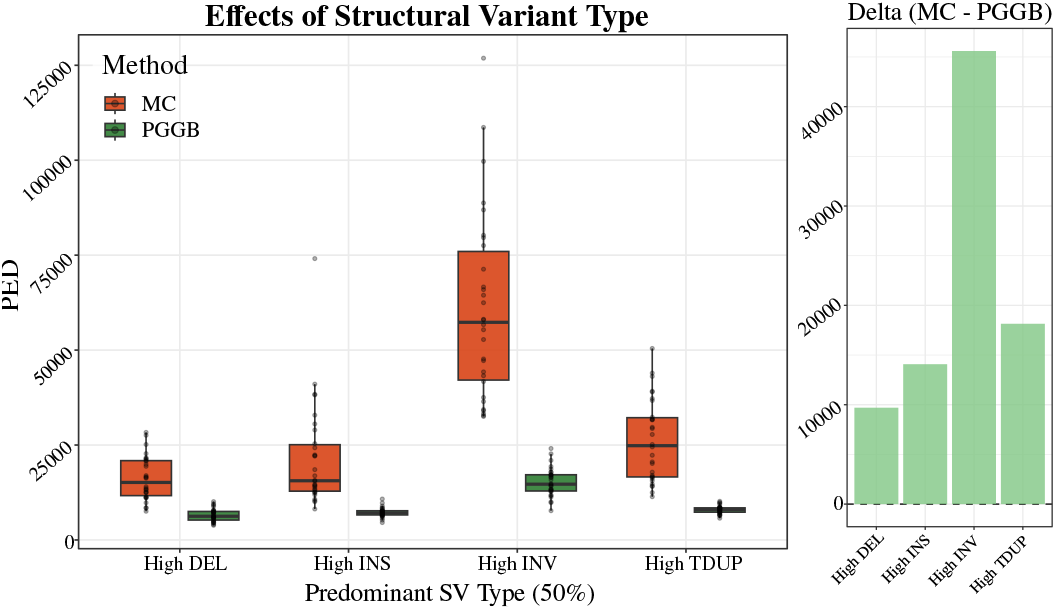
Effect of dominant variant type on reconstruction quality. Each experimental condition enriches one variant class to 55% of total structural variants. Graphical conventions follow Fig. 7.

Both tools exhibit greatest difficulty with inversions and tandem duplications. This shared limitation likely reflects the topological nature of these variant classes and the representation choices made by graph constructors: unlike insertions or deletions, which create or remove sequence, duplications and inversions are commonly represented through node reuse within the graph—repeated traversal for duplications (forming loops) and reverse-orientation traversal for inversions. Correctly encoding such structures demands accurate assessment of allelic homology, which may be hindered by alignment parameters or limitations induced by the graph constructors themselves.

To further characterize tool behavior, we quantified total node counts across all 510 graphs (Table 3). MC consistently produces substantially more nodes than `PGGB`: across all graphs, MC generates 10,997,911 nodes compared to 4,822,568 for `PGGB`—a 2.3-fold difference. This disparity is markedly amplified in inversion-enriched graphs, where MC produces 533,030 nodes versus 128,303 for `PGGB`, representing a 4.2-fold increase. These observations raise the possibility that MC’s graph representation is particularly sensitive to topologically complex variants, though the precise mechanisms underlying this inflation remain to be elucidated.

**Table 3.** Node counts across reconstruction methods.

| Condition | MC | PGGB | Ratio |
| --- | --- | --- | --- |
| All graphs ($n = 510$) | 10,997,911 | 4,822,568 | 2.28 |
| Inversion-enriched | 533,030 | 128,303 | 4.15 |

#### Joint effect of diversity and sample size

In practice, pangenome construction involves multiple parameters simultaneously, and their interaction may reveal additional insights into tool behavior. We therefore examined the joint effect of genetic diversity and sample size across the parameter space (*θ, n*) *∈ {*10^*−*3^, 10^*−*4^ *}× {* 2, 6, 12, 24*}*.

Because PED mechanically increases with haplotype count *n* (more paths require more potential edits), direct comparison across sample sizes is uninformative. We therefore introduced a normalized ratio to isolate the diversity effect at fixed sample size:

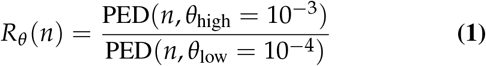

This ratio quantifies the fold-change in reconstruction error when diversity increases ten-fold, controlling for sample size effects. If tools responded identically to diversity regardless of sample size, *R*_*θ*_ (*n*) would remain constant across *n*.

The observed patterns suggest complex dynamics (Fig. 9). At low sample size (*n* = 2), the two tools exhibit markedly different sensitivity to diversity: `PGGB` shows a high ratio (*R*_*θ*_ *≈* 9.8) while MC shows a substantially lower ratio (*R*_*θ*_ *≈* 3.2). As sample size increases, these values converge, crossing around *n* = 12 where both tools show similar ratios (*R*_*θ*_ *≈* 7). At the largest sample size tested (*n* = 24), the pattern reverses: MC exhibits higher sensitivity to diversity (*R*_*θ*_ *≈* 8.8) than `PGGB` (*R*_*θ*_*≈* 7.0).

**Fig. 9.**
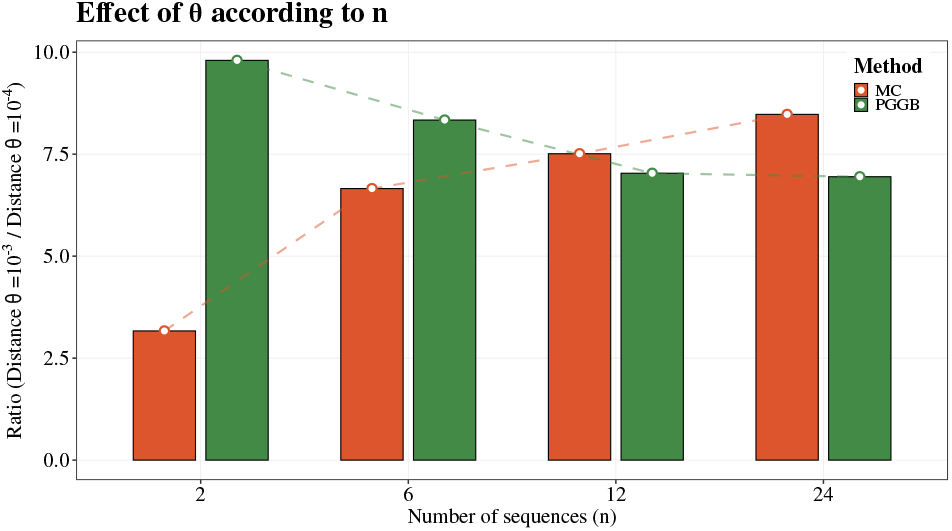
Joint effect of sample size (*n*) and genetic diversity (*θ*) on reconstruction quality (n=30). Bar height represents *R*_*θ*_ (*n*), the ratio of PED at high diversity (*θ* = 10^*−*3^) to PED at low diversity (*θ* = 10^*−*4^). Higher values indicate greater sensitivity to diversity increase. Dashed lines illustrate trends for `PGGB` (green, decreasing) and MC (orange, increasing), which cross near *n* = 12.

These crossing trajectories hint at distinct scaling behaviors between tools, though interpretation requires caution given the limited parameter space explored. One possible explanation is that `PGGB`’s all-versus-all approach benefits from additional genomes through increased information redundancy, potentially enabling more robust alignment consensus as *n* grows. Conversely, MC’s iterative, reference-guided strategy may accumulate alignment ambiguities as genomes are added sequentially, particularly under high diversity conditions. However, these mechanistic interpretations remain speculative and would require targeted experiments—such as analysis of intermediate alignment states or systematic parameter sweeps—to confirm. Moreover, the observed differences may arise not only from the underlying alignment strategies but also from heuristic choices implemented throughout the graph construction pipelines, whose effects are not always explicitly documented and are therefore difficult to assess independently.

## Discussion

`MSpangenome` introduces a genealogy-aware framework for simulating pangenomes from explicit demographic and evolutionary scenarios, generating graph structures that naturally incorporate recombination, incomplete lineage sorting and allow complex nested structural variation. Beyond producing biologically plausible synthetic datasets, this approach addresses a major methodological bottleneck in pangenomics: the lack of ground-truth reference graphs for rigorous evaluation of construction methods. The benchmarking results presented here provide an initial validation of this framework, while also highlighting the broader potential of genealogy-based pangenome simulations for building tool usage recommendations, accelerating methodological developments and testing biological hypotheses.

### Benchmarking insights: PGGB versus MinigraphCactus

Our comparison of `PGGB` and MC against a known reference reveals distinct behavioral regimes that are broadly consistent with the known algorithmic properties of both tools, lending credibility to our simulation-based evaluation framework. Under low-to-moderate genetic diversity (*≤ θ* 2 *×* 10^*−*3^), both tools achieve comparable reconstruction fidelity, with `PGGB` exhibiting marginally lower PED. The sharp transition observed at *θ* = 10^*−*2^, where PED increases by approximately two orders of magnitude, likely marks the saturation limit of alignment heuristics beyond which mutation density overwhelms the signal used to establish homology. Notably, MC outperforms `PGGB` at this extreme diversity level, suggesting that reference-guided anchoring provides more robust support than all-versus-all alignment under conditions of high sequence divergence. However, the substantially higher variance associated with MC’s performance at this threshold indicates that reconstruction stability remains precarious at our baseline diversity level (10^*−*4^).

The analysis of variant-type sensitivity exposes a shared limitation of both tools: difficulty in reconstructing inversions and tandem duplications. Unlike insertions or deletions, which create or remove sequence, these variant classes require node reuse within the graph—repeated traversal for duplications and reverse-orientation traversal for inversions. When alignment parameters or intrinsic aligner limitations prevent correct assessment of allelic homology, the alternative allele is erroneously encoded. This inflates graph complexity and PED. The quantification of total node counts across all 510 graphs corroborates this interpretation: MC produces 2.3-fold more nodes than `PGGB` overall, rising to 4.2-fold in inversion-enriched graphs. These observations highlight a persistent challenge in the field: although pangenomes are globally sparse and quasi-linear, current alignment-based methods still struggle to reconstruct the locally complex topologies created by nested genomic rearrangements and other highly repetitive or structurally complex regions.

The joint analysis of diversity and sample size further differentiates the two tools. The crossing trajectories of *R*_*θ*_ (*n*) suggest that `PGGB`’s all-versus-all strategy benefits from additional genomes through increased information redundancy, enabling more robust alignment consensus as *n* grows. Conversely, MC’s iterative, reference-guided approach appears to accumulate alignment ambiguities as genomes are added sequentially under high-diversity conditions. These mechanistic interpretations, while consistent with the algorithmic designs of both tools, remain speculative and would require targeted experiments—such as analysis of intermediate alignment states or systematic parameter sweeps—to confirm.

### The ground-truth assumption

Our benchmarking framework relies on the assumption that simulated graphs constitute an ideal topological representation of the pangenome. While this assumption is reasonable within our controlled framework, it remains partially arbitrary. Pangenomics currently lacks a universally agreed-upon standard of representation: for a given dataset, multiple graph topologies may be considered biologically valid depending on modeling choices such as node granularity or treatment of repetitive regions. This absence of consensus complicates the interpretation of PED values in absolute terms, though relative comparisons between tools remain meaningful (Bocs et al., 2025).

Furthermore, PED, while effective for quantifying global reconstruction error, does not finely characterize local topological differences. Because structural breakpoints manifest in pangenome graphs as novel edges bypassing the linear backbone, complementary metrics, such as a *breakpoint divergence ratio* measuring the proportion of unshared junction points between simulated and reconstructed graphs, would provide a normalized and more granular assessment of reconstruction quality.

### Comparison with related work

The need for controlled benchmarking data in pangenomics is underscored by the recent publication of a workflow pursuing a similar objective (Kopalli et al., 2025). However, that approach presents several limitations that `MSpangenome` was specifically designed to overcome: it lacks explicit modeling of recombination, supports a restricted spectrum of structural variants (notably excluding inversions), and relies on existing graph construction tools to generate the reference PVG. This last point complicates interpretations, as it becomes difficult to determine whether observed differences between the reference and reconstructed graphs originate from errors in the reference itself or in the reconstruction. `MSpangenome`’s *de novo* graph construction algorithm, operating directly from simulated genealogies without intermediate alignment or assembly steps, circumvents this circular dependency entirely.

### Current limitations and future directions

A fundamental constraint of `MSpangenome` stems from the coalescent-with-recombination formalism upon which demographic simulations are based. In `msprime`, the partitioning of the genome into recombination blocks (loci) is determined prior to genealogy computation, and these block boundaries remain fixed throughout the simulation. Consequently, structural variants cannot span multiple loci at any given time point, although such events are biologically plausible. In the current implementation, variant extent is therefore bounded by the interval [*i, j*] of the enclosing locus, which can become restrictively short under high population-scaled recombination rates (*ρ*)—a regime commonly observed in species such as *Drosophila melanogaster, Mytilus* spp., *Heliconius* spp., or *Quercus* spp.

Several avenues for improvement have been identified for future versions. First, enabling variants to span recombination breakpoints would allow more realistic modeling of large-scale rearrangements. Second, incorporating heterogeneous mutation rate maps along the genome would improve biological realism by simulating conserved regions (e.g., coding sequences) and mutational hotspots (e.g., telomeres, centromeres). Third, integrating transposable element libraries for generating insertions would better capture the dynamics of these pervasive structural variants, which are particularly abundant in plant genomes. Fourth, adding support for inter-chromosomal translocations would expand the spectrum of simulated rearrangements, as chromosomes are currently treated independently. Fifth, the coordinate propagation mechanism underlying nested variant generation incurs *O* (*m*^2^) worst-case complexity per locus, which may become limiting for very large sample sizes. For simulations exceeding approximately one thousand individuals, algorithmic improvements—potentially leveraging dynamic programming—will be required to maintain tractable runtimes. Finally, the inherent computational overhead of the current Object-Oriented Python implementation could be significantly mitigated by porting the core engine to a compiled systems language such as Rust.

## Conclusion

We have presented `MSpangenome`, a genealogy-aware framework for simulating pangenomes from explicit demographic and evolutionary scenarios. By coupling coalescent-based ancestry simulation with *de novo* graph construction, the framework generates pangenome variation graphs whose evolutionary origin and topology are known by construction, providing a unique source of ground-truth data for methodological evaluation.

Using these controlled references, we benchmarked two widely used graph construction tools, `PGGB` and `MinigraphCactus`, revealing contrasting strengths and limitations across diversity regimes, sample sizes, and classes of structural variation. More broadly, these results demonstrate the value of simulation-based benchmarking for identifying reconstruction errors and algorithmic failure modes that remain difficult to detect using empirical datasets alone.

Beyond benchmarking, the genealogy-aware design of `MSpangenome` opens new opportunities for investigating how demographic history, recombination, and structural variation jointly shape pangenome architecture. We anticipate that the framework will contribute both to the development of more robust pangenomics methods and to the investigation of the impact of evolutionary processes on pangenome graph structure.

## Data availability

The `MSpangenome` source code, documentation, and container recipes are publicly available at https://forge.inrae.fr/pangepop/MSpangenome. Scripts, configuration files, and analysis notebooks used for the `PGGB`/`MinigraphCactus` benchmarking are available at https://github.com/Lucien-Piat/MSpangenome_methods. Analyses were performed using MSpangepop v0.1.3.

## ACKNOWLEDGEMENTS

We thank the AgroDiv and GET-A-PAN (pangenomes.mathnum.inrae.fr) pangenome working groups, as well as the participants of the GraPanPhy and PANANNOT projects, for helpful discussions, feedback, and contributions to several pangenome variation graph workshops and hackathons. Valuable comments on the manuscript were provided by Christophe Klopp, Claire Lemaitre, Camille Marchet and Jean Monlong. L.P. and S.De. were funded by two INRAE apprenticeship grants. S.Du. was supported by state funding managed by the French National Research Agency under the France 2030 program [grant number ANR-22-PEAE-0005]. The GET-A-PAN network is supported by the INRAE departments BAP, ECODIV, GA, MathNum, MICA and SPE. The AgroDiv project is part of the Agroecology and Digital Technologies research program and received government funding managed by the Agence Nationale de la Recherche under the France 2030 program [ANR-22-PEAE-0005].

## Author contributions

L.P. (Conceptualization, Formal analysis [lead], Investigation [lead], Methodology [equal], Software [lead], Writing-original draft [lead], Writing-review & editing [equal]), S.De. (Conceptualization, Formal analysis, Investigation, Methodology, Software, Writingreview & editing [equal]), S.Du. (Formal analysis, Investigation, Methodology, Writing-review & editing [equal]), B.L. (Methodology, Writing-review & editing [equal]), and L.D. (Conceptualization [lead], Funding acquisition [lead], Formal analysis, Methodology [equal], Supervision [lead], Writing-original draft, Writing-review & editing [equal]).

### Use of Artificial Intelligence (AI) and AI-assisted Tools

During the preparation of this study, the authors used ChatGPT (based on OpenAI’s GPT-5.5 model) to assist with language refinement and editorial improvements, and Claude Opus 4.5 to assist with software development, including code suggestions and debugging support. No AI tools were used for data generation, analysis, interpretation, scientific hypothesis generation, experimental design, or decision-making. All AI-assisted outputs, including text and code, were critically reviewed, edited, tested, and validated by the authors, who take full responsibility for the final manuscript, the software, and the integrity of the results.

## Notes

https://forge.inrae.fr/pangepop/MSpangepop

https://github.com/Lucien-Piat/MSpangenome_methods/tree/main/scripts

https://github.com/inrae/MSpangepop

### Competing Interest Statement

The authors have declared no competing interest.

