## Supplemental Table 4-5 for "Simulating population pangenomes under coalescent demographic models with MSpangenome"

**Table 4.** MSpangepop simulation parameters organized by category. Parameters marked with † accept range specifications (`min`, `max`, `distribution`) for stochastic sampling across replicates.

| Parameter | Symbol | Description | Example |
|---|---|---|---|
| *Genomic parameters* | | | |
| Reference genome | | Compressed FASTA file serving as ancestral sequence | `genome.fa.gz` |
| Chromosome count | $C$ | Number of chromosomes to simulate | 12 |
| Random seed | | Seed for reproducibility (optional) | 42 |
| *Evolutionary parameters* | | | |
| Mutation rate† | $\mu$ | Per-base, per-generation mutation probability | $10^{-8}$ |
| Recombination rate† | $r$ | Per-base, per-generation recombination probability | $10^{-8}$ |
| Generation time | $g$ | Years per generation (optional, for reference) | 25 |
| *Structural variant parameters* | | | |
| SV distribution† | $P_\tau$ | Relative proportion of each variant type (%) | SNP:50, DEL:20 |
| Maximum SV count† | $V_{\max}$ | Upper bound on variant count per type (optional) | INV:20 |
| SV length distribution | | Custom TSV files specifying length distributions | `del_lengths.tsv` |
| Minimum SV length | $k_{\min}$ | Minimum variant size in base pairs | 1 |
| *Population structure parameters* | | | |
| Population ID | | Unique identifier for each population | `pop1` |
| Effective size† | $N_e$ | Effective population size | 5,000 |
| Sample size | $n$ | Number of haploid individuals sampled per population | 10 |
| Migration matrix | $\mathbf{M}$ | Pairwise migration rates between populations | |
| *Demographic event parameters* | | | |
| Event time† | $T$ | Time of event in generations (backward) | 3,000 |
| Size change† | $N_e'$ | New effective size after population size change | 10,000 |
| Migration rate change† | $m$ | New migration rate between specified populations | $10^{-4}$ |
| Mass migration proportion† | $p_m$ | Fraction of lineages migrating instantaneously | 0.2 |
| *Workflow parameters* | | | |
| Output directory | | Base path for simulation results | `results/` |
| Replicates | $R$ | Number of independent simulation runs | 100 |
| Memory multiplier | | Scaling factor for cluster memory allocation | 1.5 |
| Succinct mode | | Skip visualization rules if `True` | `False` |

**Table 5.** Supported demographic events in MSpangepop, following `msprime` conventions (?).

| Event type | Description |
|---|---|
| `population_parameters_change` | Instantaneous change in $N_e$ or growth rate |
| `add_migration_rate_change` | Modify continuous migration rate between populations |
| `mass_migration` | Instantaneous movement of lineage fraction between populations |
| `population_split` | Division of ancestral population (forward in time) |
| `admixture` | Merging of lineages from multiple source populations |
